# Inoculum size effects in *bla*_*IMP-6*_ and plasmid-mediated quinolone resistance gene-positive Enterobacteriaceae in Japan

**DOI:** 10.1101/534958

**Authors:** Yoshihiko Ogawa, Ryuichi Nakano, Kei Kasahara, Tomoki Mizuno, Nobuyasu Hirai, Akiyo Nakano, Yuki Suzuki, Naoki Kakuta, Takashi Masui, Hisakazu Yano, Keiichi Mikasa

## Abstract

The aim of this study was to examine the resistance genes in clinical isolates which produced IMP-6 type metallo-β-lactamase lactamase (MBL) and had mildly reduced susceptibilities to levofloxacin and/or amikacin. The inoculum size effect was also assessed. A total of 14 Enterobacteriaceae isolates (2 *Escherichia coli* and 12 *Klebsiella pneumoniae*) which produced IMP-6 MBL, and had mild increases in their MICs for levofloxacin and amikacin were examined. Thirteen out of 14 isolates harbored CTX-M-2, with the remaining isolate co-harboring CTX-M-2 and CTX-M-1 as ESBLs. All isolates carried one or more PMQRs; *aac(6′)-Ib-cr* was the most prevalent (92.8%), followed by *oqxA* (64.3%), *qnrS* (42.9%), *oqxAB* (21.4%), and *qnrB* (14.3%). The inoculum size effects were significant in all strains for meropenem, 13 for imipenem, 7 for levofloxacin, and 3 for amikacin. Conjugation was successfully performed with 8 isolates and 11 strains were obtained. Eleven of the experimental strains (100%), and 8 strains (72.7%) showed inoculum size effects for meropenem and imipenem, respectively. No inoculum size effect was seen for levofloxacin. Four strains harbored *qnr* genes and 2 strains harbored *qnr* genes and QRDR mutations concurrently. *bla*_IMP-6_ positive Enterobacteriaceae with mildly reduced susceptibilities to levofloxacin and/or amikacin also harbored at least one plasmid-mediated drug resistance gene. These represent an unrecognized threat, capable of compromising the *in vitro* activity of many classes of antimicrobial agents. We conclude that IMP-6 MBL plays an important role in decreasing the MIC for carbapenems, whereas *qnr* does not for levofloxacin.

## Introduction

Antimicrobial resistance in Gram-negative bacteria is an emerging and serious global public health threat. Most importantly, carbapenemase-producing Enterobacteriaceae (CPE) have emerged, which confer broad resistance to most β-lactam antibiotics including the “last-resort” carbapenems (1-3). Although the number of CPE infection cases is increasing, the optimal treatment paradigm for CPE infections has not been defined. Furthermore, there are numerous different types of carbapenemase enzymes, each conferring varying spectrums of resistance.

Much of the existing knowledge arises from reviews of case series and retrospective studies on VIM- or (Klebsiella pneumoniae carbapenemase) KPC-producing CPEs which are now wide-spread in Europe and the United States. Based on these published data, combination therapy is recommended for CPE infections in reference to the results of drug susceptibility tests. Most cases of CPE infection are caused by *bla*_IMP_ positive Gram-negative bacteria (especially *bla*_IMP-6_) in Japan (4) and these organisms show a susceptibility to imipenem (5). Because of the good susceptibilities at a glance, passing over of the isolates with this resistant gene leaded to the outbreaks. On the other hand, fortunately, many of the *bla*_IMP-6_ positive CPEs are susceptible to levofloxacin and amikacin (6). Thus, based on the results of drug susceptibility tests, some infectious cases caused by IMP-type carbapenemase producing Enterobacteriaceae have been treated with quinolone monotherapy (7).

Quinolone resistance in Gram-negative bacteria is mostly mediated by point mutations that arise in the quinolone resistance-determining regions (QRDRs) of the gyrase and topoisomerase IV genes, leading to a modification of the target (8). However, the previously unidentified resistance to quinolones mediated by plasmid-mediated quinolone resistance genes (PMQRs) has also begun to be recognized as a drug resistance mechanism (8-10). These genes were first identified in 1998 and include the qnr proteins, aminoglycoside acetyltransferase, and the plasmid-mediated efflux pumps QepA and OqxAB (10). *Aac(6′)-Ib-cr* is one variant of aminoglycoside acetyltransferase which can reduce the activity of ciplofloxacin and aminoglycosides, including amikacin (8).

The inoculum size effect is a phenomenon whereby the measured value of the MIC changes depending on the number of bacteria, and its effect on β-lactamases (including metallo-β-lactamases: MBLs) has been described (11-12). More recently, the inoculum size effect for the action of quinolone on bacteria expressing PMQR genes has also been described (9,13). However, in one of these studies, it was reported that only half of the transconjugants of PMQRs (2/4 strains) showed an inoculum size effect for quinolone (9). There has been no study describing the inoculum size effect for strains expressing both MBL and PMQRs.

The aim of this study was to examine the drug resistance genes found in clinical isolates which produce IMP-6 type carbapenemase and have mildly reduced susceptibilities to levofloxacin and/or amikacin. In addition, we assessed changes in the MICs for antibiotics and the impact of these changes.

## Methods

### Bacterial isolates

Sequentially collected clinical isolates of *bla*_IMP-6_ positive Enterobacteriaceae from Japan, which showed a mild decrease in susceptibility to levofloxacin and/or amikacin (the range of MICs for levofloxacin was 0.5–4 μg/mL and for amikacin it was 2–16 μg/mL, determined using the agar dilution method according to CLSI guidelines M100-S25) (14), were studied and included *Escherichia coli* (n = 2) and *Klebsiella pneumoniae* (n = 12). These isolates were non-duplicated and were obtained from 12 hospitals.

### Detection of antimicrobial resistance genes

All isolates were confirmed *bla*_IMP-6_ positive by both PCR and DNA sequencing (15), and we performed additional PCR testing for *bla*_VIM_, *bla*_KPC_, *bla*_NDM_, and *bla*_OXA-48-like_ to assess additional carbapenemase genes (16). We also performed PCR to detect the presence of extended-spectrum β-lactamases (ESBLs; TEM, SHV, CTX-M, and OXA) (17). The presence of plasmid-mediated quinolone resistance genes (*qnrA, qnrB, qnrC, qnrD,* and *qnrS*), efflux pump genes (*qepA* and *oqxAB*), *aac(6′)-Ib* and *aac(6′)-Ib-cr* was also assessed by PCR (18-21).

The presence of aminoglycoside-modifying enzymes (*aph(3′)-VI* and *ant(4′′)-IIa*), which have been reported to reduce susceptibility to amikacin and have been detected in Enterobacteriaceae, was also assessed by PCR using specific primers (22,23). The QRDRs in *gyrA* and *parC* were amplified as previously described (24,25) and were sequenced to assess any co-existing chromosomal mutations (26).

### Antimicrobial susceptibility testing

The minimum inhibitory concentrations (MICs) of levofloxacin, amikacin, imipenem, and meropenem, were evaluated by the agar dilution method at a standard inoculum size according to the CLSI guidelines (27). We also evaluated the MICs of these drugs for the strains using 10- and 100-fold of the colony forming units (CFU) in the standard inoculum. All results were interpreted according to the CLSI criteria describing *in vitro* susceptibility (14). We defined that an inoculum size effect was significant if the MICs of antibiotics showed ≥ 4-fold increase at a 100-fold of the inoculum size compared with the standard inoculum size.

### Conjugation experiments

Conjugation experiments were performed using the broth mating technique with sodium azide resistant *E. coli* J53 and *E. coli* NR 3500 containing the gyrA mutation Ser83Phe. *E. coli* NR3500 were obtained from *E. coli* J53 using an LB agar plate containing levofloxacin (MIC; 0.125 μg/mL). Transconjugants were selected on LB agar plates containing sodium azide (100 μg/mL), and cefpodoxime (8 μg/mL). Transfer of drug resistant genes (IMP-6, CTX-M-1, CTX-M-2, *qnrB, qnrS, oqxA, oqxB*, and *aac(6′)Ib-cr*) was confirmed by the PCR method described above.

## Results

### Antibiotic resistant genes

For carbapenemase genes, none of the tested isolates harbored other MBL genes besides *bla*_IMP-6_. For ESBL genes, CTX-M-2 was detected in all isolates, and one isolate co-harbored both CTX-M-2 and CTX-M-1. For PMQR encoding genes, 13 isolates (92.8%) harbored *aac(6′)-Ib-cr,* followed by *oqxA* (9 isolates, 64.3%), *qnrS* (6 isolates, 42.9%), *oqxAB* (3 isolates, 21.4%) and *qnrB* (2 isolates, 14.3%) (Table 1). *oqxA* and *B* were not detected in *E. coli* nor in any of the transconjugants. None of the tested isolates harbored *qnrA, C, D*, or *qepA*. In addition, none of the isolates harbored *aph(3′)VI* and *ant(4′′)IIa* as the aminoglycoside modifying enzyme genes that have been reported to reduce susceptibility to amikacin (Table 1).

**Table 1.** Phenotypic and genotypic characteristics of clinical isolates of *bla*_IMP-6_ positive Enterobacteriaceae

| Isolates | Organism | Resistance genes | *gyrA* mutation | *parC* mutation | MIC (μg/mL) |  |  |  |  |  |  |  |  |  |  |  |
| --- | --- | --- | --- | --- | --- | --- | --- | --- | --- | --- | --- | --- | --- | --- | --- | --- |
|  |  |  |  |  | Levofloxacin |  |  | Amikacin |  |  | Imipenem |  |  | Meropenem |  |  |
|  |  |  |  |  | Standard | × 10 | × 100 | Standard | × 10 | × 100 | Standard | × 10 | × 100 | Standard | × 10 | × 100 |
| NR286 | *E. coli* | IMP-6,CTX-M-2, *qnrS*, *aac(6')-Ib-cr* |  |  | 0.5 | 0.5 | **8** | 16 | 32 | 32 | 0.125 | 0.25 | **1** | 2 | 8 | **16** |
| NR379 | *E. coli* | IMP-6, CTX-M-2, *qnrB*, *aac(6')-Ib-cr* | Ser83Leu |  | 0.5 | 2 | **8** | 8 | 16 | **32** | 0.5 | 0.5 | 1 | 4 | 8 | **32** |
| NR411 | *K. pneumoniae* | IMP-6, CTX-M-2, *aac(6')-Ib-cr*, *oqxAB* | Asp87Asn |  | 4 | 8 | **16** | 2 | 2 | **8** | 0.125 | 0.125 | **1** | 2 | 8 | **64** |
| NR417 | *K. pneumoniae* | IMP-6, CTX-M-2, *aac(6')-Ib*, *oqxA* | Ser83Try | Ser80Ile | 1 | 2 | **4** | 4 | 8 | 8 | 0.125 | 0.25 | **2** | 1 | 8 | **64** |
| NR420 | *K. pneumoniae* | IMP-6, CTX-M-2, *qnrS*, *aac(6')-Ib-cr*, *oqxA* |  |  | 0.5 | 1 | **4** | 4 | 8 | 8 | 0.125 | 0.5 | **2** | 2 | 16 | **64** |
| NR449 | *K. pneumoniae* | IMP-6, CTX-M-2, *qnrS*, *aac(6')Ib-cr*, *oqxAB* |  |  | 1 | 1 | 2 | 2 | 2 | 2 | 0.125 | 0.25 | **4** | 1 | 8 | **64** |
| NR462 | *K. pneumoniae* | IMP-6, CTX-M-2, *aac(6')-Ib-cr*, *oqxA* | Asp87Gly |  | 1 | 1 | 1 | 2 | 2 | 2 | 0.125 | 0.25 | **0.5** | 1 | 2 | **32** |
| NR465 | *K. pneumoniae* | IMP-6, CTX-M-2, *qnrS*, *aac(6')-Ib-cr*, *oqxAB* |  |  | 1 | 2 | 2 | 2 | 2 | 4 | 0.063 | 0.125 | **2** | 1 | 16 | **64** |
| NR487 | *K. pneumoniae* | IMP-6, CTX-M-2, *qnrS*, *aac(6')-Ib-cr*, *oqxA* |  |  | 1 | 1 | **4** | 2 | 4 | 4 | 0.25 | 0.5 | **2** | 1 | 16 | **64** |
| NR490 | *K. pneumoniae* | IMP-6, CTX-M-2, *aac(6')-Ib-cr*, *oqxA* | Asp87Gly |  | 4 | 4 | 4 | 8 | 8 | 16 | 0.125 | 0.5 | **2** | 1 | 16 | **128** |
| NR496 | *K. pneumoniae* | IMP-6, CTX-M-2, *qnrS*, *aac(6')-Ib-cr*, *oqxA* |  |  | 1 | 1 | **4** | 2 | 4 | 4 | 0.125 | 0.5 | **2** | 1 | 8 | **64** |
| NR499 | *K. pneumoniae* | IMP-6, CTX-M-1, CTX-M-2, *qnrB*, *qnrS*,<br>*aac(6')-Ib-cr*, *oqxA* |  |  | 0.5 | 0.5 | 1 | 2 | 4 | **8** | 0.25 | 0.5 | **2** | 2 | 16 | **128** |
| NR512 | *K. pneumoniae* | IMP-6, CTX-M-2, *aac(6')-Ib-cr*, *oqxA* |  |  | 1 | 2 | 2 | 2 | 2 | 4 | 0.125 | 0.25 | **1** | 1 | 8 | **64** |
| NR1647 | *K. pneumoniae* | IMP-6, CTX-M-2, *aac(6')-Ib-cr*, *oqxA* |  |  | 0.5 | 0.5 | 1 | 2 | 2 | 2 | 0.25 | 0.25 | **2** | 4 | 16 | **64** |

Sequencing of the PCR products derived from the QRDRs in *gyrA* and *parC* showed a substitution in the QRDR of GyrA in 1 *E.coli* (NR379) and 4 *K. pneumoniae* isolates (NR411, 417, 462, 490), and in the QRDR of ParC (NR417).

### Drug susceptibility tests and inoculum size effects

The results of the antibiotic susceptibility tests are shown in Table 1. For all the antibiotics tested, the MIC values of all the isolates at 10-fold the standard inoculum size were equal to or higher than those using the standard inoculum, and those at 100-fold the standard inoculum size were equal to or higher than those using 10-fold the standard inoculum size. Inoculum size effects were observed in all isolates for meropenem (MIC range; 1–4 to 16–128 μg/mL), 13 for imipenem (MIC range; 0.063–0.5 to 0.5–2 μg/mL), 7 for levofloxacin (MIC range; 0.5–4 to 1–16 μg/mL), and 3 for amikacin (MIC range; 2–16 to 2–32 μg/mL). Based on the CLSI breakpoint in M100-S25 at the standard inoculum size, all isolates showed susceptibility to amikacin and imipenem. Two isolates (14.3%) and 6 isolates (42.9%) were not susceptible to levofloxacin and meropenem, respectively. On the other hand, at 100-fold the standard inoculum size, 2 isolates, 8 isolates, 9 isolates were found to not be susceptible to amikacin, levofloxacin, and imipenem, respectively. All isolates were resistant to meropenem at 100-fold the standard inoculum size.

### Conjugation experiments

Conjugation experiments were performed successfully with 8 isolates and 11 strains were obtained (Table 2). The recipient strain (*E. coli* J53, NR3500) did not show an inoculum size effect and was susceptible to all of the antibiotics we tested with or without the presence of QRDR mutation. Eleven isolates (100%), 8 strains (72.7%), and 1 strain showed inoculum size effects for meropenem, imipenem, and amikacin, respectively. In addition, based on our definition, no inoculum size effect for levofloxacin was seen in the conjugant strains. In these strains, 4 strains harbored *qnr* genes, and 2 strains harbored both *qnr* genes and QRDR mutation concurrently. Thus, our experiments showed that the inoculum size effects for carbapenems were apparent with IMP-6 type carabapenemase, and that these effects were greater for meropenem than for imipenem. In addition, the *qnrS* and *qnrB* genes did not provide an apparent inoculum size effect for levofloxacin. Similarly, the *aac(6′)Ib-cr* gene did not provide an inoculum size effect for amikacin.

**Table 2.** Phenotypic and genotypic characteristics of the transconjugants of *bl*_*a*IMP-6_ producing *E. coli*

| Strain | Resistance genes | *gyrA* mutation | MIC (µg/mL) |  |  |  |  |  |  |  |  |  |  |  |
| --- | --- | --- | --- | --- | --- | --- | --- | --- | --- | --- | --- | --- | --- | --- |
|  |  |  | Levofloxacin |  |  | Amikacin |  |  | Imipenem |  |  | Meropenem |  |  |
|  |  |  | Standard | × 10 | × 100 | Standard | × 10 | × 100 | Standard | × 10 | × 100 | Standard | × 10 | × 100 |
| J53 *E. coli* |  |  | 0.031 | 0.031 | 0.063 | 1 | 1 | 1 | 0.25 | 0.25 | 0.5 | 0.031 | 0.063 | 0.063 |
| NR3500 |  | Ser83Phe | 0.5 | 0.5 | 0.5 | 0.5 | 0.5 | 1 | 0.125 | 0.125 | 0.25 | 0.031 | 0.031 | 0.063 |
| NR286/J53 | IMP-6, CTX-M-2, *aac(6')Ib-cr* |  | 0.031 | 0.031 | 0.031 | 4 | 4 | 8 | 0.5 | 0.5 | **2** | 1 | 4 | **32** |
| NR379/J53-1 | IMP-6, CTX-M-2, *aac(6')Ib-cr* |  | 0.031 | 0.031 | 0.063 | 4 | 4 | 4 | 0.5 | 0.5 | 1 | 0.5 | 2 | **32** |
| NR379/J53-2 | IMP-6, CTX-M-2, *qnrB*, *aac(6')Ib-cr* |  | 0.031 | 0.031 | 0.031 | 16 | 16 | 16 | 0.5 | 0.5 | 1 | 0.5 | 1 | **16** |
| NR379/NR3500 | IMP-6, CTX-M-2, *qnrB* | Ser83Phe | 1 | 1 | 2 | 8 | 8 | 8 | 0.125 | 0.125 | 0.25 | 0.25 | 0.5 | **4** |
| NR420/J53 | IMP-6, CTX-M-2, *aac(6')Ib-cr* |  | 0.031 | 0.031 | 0.031 | 1 | 1 | 2 | 0.25 | 0.5 | **2** | 0.5 | 4 | **32** |
| NR449/J53 | IMP-6, CTX-M-2, *aac(6')Ib-cr* |  | 0.031 | 0.031 | 0.063 | 0.5 | 0.5 | 1 | 0.125 | 0.25 | **1** | 0.25 | 2 | **32** |
| NR462/J53 | IMP-6, CTX-M-2, *aac(6')Ib-cr* |  | 0.031 | 0.031 | 0.063 | 0.5 | 0.5 | 1 | 0.125 | 0.25 | **0.5** | 0.25 | 2 | **32** |
| NR487/J53 | IMP-6, CTX-M-2, *qnrS*, *aac(6')Ib-cr* |  | 0.031 | 0.031 | 0.063 | 1 | 1 | 2 | 0.5 | 0.5 | **2** | 0.5 | 4 | **32** |
| NR487/NR3500 | IMP-6, CTX-M-2, *qnrS* | Ser83Phe | 0.5 | 0.5 | 0.5 | 0.5 | 1 | **2** | 0.125 | 0.25 | **1** | 0.25 | 1 | **16** |
| NR499/J53 | IMP-6, CTX-M-2, *aac(6')Ib-cr* |  | 0.031 | 0.031 | 0.031 | 1 | 1 | 2 | 0.25 | 0.5 | **2** | 0.25 | 1 | **32** |
| NR512/J53 | IMP-6, CTX-M-2, *aac(6')Ib-cr* |  | 0.031 | 0.031 | 0.031 | 1 | 1 | 1 | 0.25 | 0.5 | **1** | 0.25 | 4 | **4** |

## Discussion

*bla*_IMP_ positive Enterobacteriaceae have recently become a serious problem throughout Asian countries, including Japan (4). However, there are very few studies that have reported other drug resistance genes besides β-lactamases in Japan. We have found that *bla*_IMP-6_ positive Enterobactericeae with moderately increased MICs for levofloxacin and amikacin harbor multiple drug resistant genes including ESBLs and PMQRs. To the best of our knowledge, this is the first report evaluating drug resistance genes other than β-lactamases and the inoculum size effects for isolates producing IMP-6 type MBL.

The results of our conjugation experiments are in line with the current knowledge that PMQRs provide only a low level of resistance to levofloxacin or amikacin. Moreover, although inoculum size effects for levofloxacin, amikacin, and carbapenems were apparent for some isolates in our study, the inoculum size effects for levofloxacin and amikacin were not significant in the conjugation experiments. Margaritis et al. have reported that half of the transconjugants of PMQRs (2/4 strains) showed inoculum size effects for levofloxacin (1-, 2-, 4-, and 16-fold: MIC of 10^7^ CFU/mL per MIC of 10^5^ CFU/mL) (9). However, they also showed that the inoculum size effects for quinolone were significant for laboratory strains which did not harbor acquired quinolone resistance genes. Thus, *qnr* genes have only a small impact on the inoculum size effects for levofloxacin. Rice reported that resistance to quinolone arises as the result of a combination of various resistance mechanisms, and that the acquisition of *qnr* genes increase the MIC but not to a level that would be considered clinically resistant (28). Further studies should be performed to identify other factors that could explain this difference.

One experimental strain (NR 379 and NR 3500), which harbored the *qnrB* gene and *gyrA* mutation, showed susceptibility to levofloxacin based on CLSI breakpoint M100-S25, but was resistant based on the EUCAST clinical breakpoint ver. 8.1 at 100-fold the inoculum size (29). Therefore, we should not completely ignore *qnr* genes if the strain shows a decreased susceptibility to levofloxacin in the setting of a huge level of organismal infection (e.g. an abscess or in bacteremia).

Presently, a combination therapeutic strategy is recommended for carbapenemase-producing Enterobacteriaceae infections (1-2, 30). In addition, Morrill et al. have recommended that an empiric treatment for carbapenem-resistant KPC-producing *K. pneumoniae* might be high-dose carbapenem and an adjunct drug such as aminoglycoside if the MIC for carbapenem was ≤ 16 μg/mL (3). These recommendations are based on clinical experiences with VIM- or KPC-type carbapenemase producing Enterobacteriaceae infections where the MICs for carbapenem are high. On the other hand, the reported susceptibility to meropenem and imipenem among *bla*_IMP-6_ positive *E. coli* was approximately 70% and 100% (4). Pang et al. have reported that most pathogens have been confirmed to produce IMP-type carbapenemases, and some cases have been successfully treated with quinolone monotherapy as a definitive therapy based on drug susceptibility tests (7). However, in the present study the isolates did not show resistance to levofloxacin, amikacin, and carbapenems based on the CLSI definition, and the MICs of some isolates turned to be high enough to be resistant to these drugs especially meropenem (the MIC for all isolates was ≥ 16 μg/mL) at 100-fold the standard inoculum size. This suggests that in the context of a clinical infection, these pathogens may fail to be treated despite the use of “susceptible” drugs identified by the usual methods. In fact, it has been reported that 55.6% of infectious cases caused by KPC-type carbapenemase producing *K. pneumoniae,* which showed susceptibility in automated drug susceptibility tests, failed to be treated by imipenem or meropenem (31).

In conclusion, *bla*_IMP-6_ positive Enterobacteriaceae, with a mild MIC elevation for levofloxacin and/or amikacin, harbored at least one plasmid-mediated drug resistance gene, other than ESBLs, and carbapenemase at the same time. Furthermore, these isolates showed inoculum size effects for levofloxacin, amikacin, and carbapenems (especially for meropenem compared with imipenem). The decrease in the MICs for carbapenems was thought to be mainly due to IMP-6 type metallo-β-lactamase, however *qnr* had a small effect on the decrease in the MIC for levofloxacin.

